# Standards needed? An Exploration of Qualifying Exams from a Literature Review and Website Analysis of University-Wide Policies

**DOI:** 10.1101/2022.11.14.516505

**Authors:** Jacqueline E. McLaughlin, Kathryn Morbitzer, Margaux Meilhac, Natalie Poupart, Rebekah L. Layton, Michael B. Jarstfer

## Abstract

**Purpose:** While known by many names, qualifying exams function as gatekeepers to graduate student advancement to PhD Candidacy, yet there has been little formal study on best qualifying exam practices particularly in biomedical and related STEM PhD programs. The purpose of this study was to examine the current state of qualifying exams through an examination of the literation and exploration of university policies.

**Design/methodology/approach:** We conducted a literature review of studies on qualifying exams and completed an external evaluation of peer institutions’ and internal institutional qualifying exam requirements to inform our discussion of common practices of qualifying exams in doctoral training at a research-intensive institutions.

**Findings:** Our research identified the need for more evidence-based research on qualifying exams, which are nearly universal as a major doctoral training milestone across US institutions of higher education. Our findings indicate a wide variety of qualifying exam formats, often lacking the establishment of explicit expectations and evidence for specific formats. This lack of understanding of best practices coupled with insufficient clarity has a real potential to disadvantage doctoral students, particularly first generation, underrepresented minority, international, and/or other trainees who are not privileged or socialized to navigate training environments with vague landmarks such as the qualifying exams.

**Originality:** There are very few studies that evaluate qualifying exams in US doctoral education, particularly in STEM fields, and to our knowledge, there has been no analysis of campus wide policies on qualifying exams reported. The lack of evidence for best practices and the need for to evaluate the implementation and effectiveness of qualifying exams are discussed.

## INTRODUCTION

The Doctor of Philosophy (PhD) is the highest academic degree for STEM (Science, Technology Engineering, and Mathematics) fields and requires completion of an intense, mentored research project. The most common structure for United States (U.S.) doctoral programs includes four elements : 1–2 years of core and elective coursework in the discipline; a qualifying exam (QE); in the form of a dissertation (Goldman and Massy, 2000; Hartnett and Katz, 1977; National Academies of Sciences, Engineering, and Medicine, 2018; Walker et al., 2008). While the specifics of each element vary by discipline and institution, they generally exist in some form in most programs (Austin, 2002; Hartnett and Katz, 1977).

Increasingly, careers of young PhD scientists are diverging from their thesis work, leading to calls for ensuring that core competencies of PhD programs include translatable skills (e.g., leadership, communication, management) in addition to technical and discipline specific competencies (e.g., scientific knowledge, technical skills, data interpretation) (McLaughlin et al., 2019; Sinche et al., 2017). Since PhD training programs are not typically accredited by a professional organization in contrast to many professional degree programs, there is no common list of competencies or core outcomes defined. Instead, each program is tasked with defining their own core competencies and program outcomes. The 2018 report from the National Academies of Sciences, Engineering, and Medicine Committee, *Revitalizing Graduate STEM Education for the 21st Century*, identified two overarching competencies: “Develop Scientific and Technological Literacy and Conduct Original Research” and “Develop Leadership, Communication, and Professional Competencies” (National Academies of Sciences, Engineering, and Medicine, 2018). Despite this attempt at defining common competencies, it remains unclear if or how these have been adopted by PhD programs and the extent to which PhD programs have defined their own core competencies – and corresponding assessments.

Our research team is pursuing a broad research program exploring the alignment of PhD core competencies, assessments, and training requirements with students’ needs. In this essay, we focus specifically on exploring how US doctoral training programs consider and utilize qualifying exams (QEs) during the course of PhD training. Although the terminology used to refer to QEs varies widely by discipline, department, and university (e.g., candidacy, comprehensive, cumulative, general, preliminary, or qualifying exam), we will refer to these as QE throughout. There is notable variability in QE formats but consistently, QEs are executed as high stakes assessments typically administered midway through a PhD program after coursework is complete to assess readiness for dissertation research. Since these are typically high stakes exams, it seems particularly critical that QEs align with core competencies defined by PhD programs. Here, we explore the current state of QEs and address specific questions around the purpose, perceptions, and format of QEs from a survey of the peer-reviewed literature, publications in various academic press sources, and from website analysis of peer institutes. Our investigation exposes a research gap concerning the purpose, format, and effectiveness of STEM QEs; reveals a lack of QE consistency across PhD programs; and identifies the need for robust investigation of QEs and their role as high-stakes exam used to evaluate PhD students’ advancement to candidacy.

## METHODS

To investigate evidence for best practices, we conducted a literature review. The literature review was initially focused on biomedical and basic science PhD programs. However, the paucity of reports on qualifying exams led us to expand to any publication that discussed qualifying and related exams in doctorate education with a focus on most recent publications. A search of the literature review was conducted by using scholarly databases, including ERIC and PubMed, with a combination of search terms including “qualifying exam”, “doctoral exam”, “preliminary exam”, “candidacy exam”, “comprehensive exam”, and “cumulative exam”.

To examine current policies, we reviewed graduate school websites. We chose programs that are considered peers of the University of North Carolina and peers of the UNC Eshleman School of Pharmacy. We examined the websites of the following universities: Duke University, Emory University, Johns Hopkins University, Northwestern University, The Ohio State University, University of Arizona, University of California-Berkeley, University of California-Los Angeles, University of California-San Francisco, University of Florida, University of Illinois at Urbana-Champaign, University of Illinois-Chicago, University of Kentucky, University of Michigan-Ann Arbor, University of Minnesota, University of North Carolina at Chapel Hill, University of Pittsburgh-Pittsburgh, University of Texas-Austin, University of Virginia-Main Campus, University of Washington-Seattle, University of Wisconsin-Madison, University of Kansas, Vanderbilt University. We manually examined graduate school websites for examination milestones required before the thesis defense and compared the examination descriptions, characteristics, and regulations.

### QUALIFYING EXAMS: LITERATURE REVIEW

Globally, PhD training involves a high stakes assessment before “candidacy”, which we will consider as the time when the only remaining formal requirement is completing the dissertation research, preparing the dissertation, and defending the thesis. Most U.S. and Canadian PhD programs include a QE to evaluate students before advancing to candidacy whereas most European and Australian PhD programs require a master’s degree for entry into PhD programs in lieu of QEs (Barnett et al., 2017; Bernstein et al., 2014). Despite their importance in U.S. PhD training, there is little peer-reviewed literature about QEs, a fact highlighted in publications starting in the 1980s, including continued calls for focused research on QEs, to the present. As a result, in this literature review, we included all publications addressing doctoral training – including various doctoral degrees - starting from the 1960s to gain a full picture of prior work. Notably, there were very few papers that considered QEs in a biomedical or adjacent discipline PhD program (e.g., pharmaceutical sciences, biology, chemistry, biochemistry, biophysics, genetics, biomedical engineering) (Wilson et al., 2018) or provided in-depth evaluation of modern QE practices in any discipline with a doctorate.

#### QE history

To understand where we currently are using QEs in PhD training, we explored the history of QEs. Doctorate programs are rarely subject to strict regulations from accrediting bodies so common practices appear to be a matter of historic precedence, acceptance of previous practices, and periodic change after internal and or external reviews. Additionally, faculty within doctoral programs may tend to be more research focused rather than teaching focused, leading to less priority being placed on the pedagogy and assessments associated with the doctoral program and more emphasis on the research performance. Prior literature reviews have noted that QEs were part of the first PhDs awarded in the United States dating as far back as the late 1800s. When PhD training expanded in the 1930s and again in the 1960s, Universities considered ways to standardized PHD training, and QEs were incorporated as part of the standard progression in most if not all PhD programs. With the expansion of PhD training came the burdens associated with assessing PhD trainees; as a result, the 1960s reforms to streamline QEs and move them earlier in training were commonly accepted (Estrem and Lucas, 2003). These changes led to the current cadence of course work, QEs, dissertation research and final defense as nearly universal in PhD training (Bernstein et al., 2014; Estrem and Lucas, 2003; Stewart ▫ Wells and Keenan, 2020).

#### The purpose of QEs

Since QEs are a shared practice, one anticipates that programs might also have a shared purpose in utilizing QEs to evaluate trainees. We found that many purposes for QEs have been expressed through the years, and these continue to evolve as doctoral programs adapt to training needs. An early 1980s survey of the literature (Anderson et al., 1984) found that QEs were intended “to screen out students on the basis of ability and/or knowledge; to provide a rite of passage so the student will feel the degree has been earned; to provide an opportunity for students to organize their thinking and integrate what has been learned”(Anderson et al., 1984). An analysis of Composition and Rhetoric PhD programs from the early 2000s (Estrem and Lucas, 2003) found that programs listed over 40 stated purposes for QEs and these could be binned into three categories: critically thinking, expert knowledge, and research/teaching ability. In a survey of Counseling Psychology Training PHD programs (Manus et al., 1992), 78% of programs reported that the purpose of QEs was to assess students’ synthesis and integration of knowledge as well as basic skills and abilities (Manus et al., 1992). Similarly, a survey of directors of social work doctoral programs (Furstenberg & Nichols▫Casebolt, 2001) found that QEs function to test “students’ mastery of knowledge and skills and assisting students’ learning and progress toward the dissertation” (Furstenberg and Nichols▫Casebolt, 2001). In addition to evaluating students, student performance on QEs can also be used for program evaluation (Cobia et al., 2005). For example, if QEs are designed around a program’s stated competencies and outcomes, demonstrated mastery of QEs can assess whether students’ training prior to the QE is meeting the trainees’ needs.

A more recent survey of faculty in doctoral counselor education programs identified five QE purposes, ranked by participants as: 1) to assess lower levels of cognitive complexity, 2) to assess higher levels of cognitive complexity, 3) to prepare student for future scholarship, 4) to promote a beneficial learning experience, and 5) to maintain tradition (Kostohryz, 2016). Other doctoral programs report similar purposes for QEs, including Biomedical Sciences (Wilson et al., 2018), criminal justice (Pelfrey and Hague, 2000), marketing (Ponder et al., 2004), political science (McMahon et al., 2020), mathematics (Earl▫Novell, 2006), and nursing (Mawn and Goldberg, 2012). Student perspectives include a perception that QEs are intended to “test the capability of the candidate and to weed out weak students,” though this finding was based on a small sample (n = 35) of engineering students (Memarian et al., 2019). Clearly, the stated and functional purposes of QEs are varied and multifaceted. If the intended purpose is not specifically stated and socialized with students, the vagueness creates a risk that students and faculty will have different perspectives of QEs. A recent but very small study of faculty and students in a Doctor of Education (EdD) program showed a split on the purpose of QEs as it related to students’ futures, with faculty viewing the QE as important for careers in academia, whereas students saw them as important as preparation for thesis work (Guloy et al., 2020).

The complex expressions of the purposes for QEs are consistent with our interpretation leading to one possible concrete, encompassing, and shared purpose of QEs: to ensure students are prepared for their dissertation research enroute to their doctorate. The fact that a transition to “candidacy” is frequently associated with passing QEs is consistent with this interpretation. Despite a complexity of purposes, the functional necessity of QEs can be distilled to ensuring that students are ready to progress to a focus on dissertation research. The fact that QEs are considered high-stake assessments should lead administrators to consider many questions when establishing QE criteria and expectations. While the following are not exclusive, we consider them to be some of the most pressing in determining the format and administration of QEs within a program:

- When should QEs be administered?
- How should QEs be administered?
- How should success of QEs be assessed?
- What is the student experience with QEs?
- What is the faculty experience with QEs?
- Are QE practices equitable? A
- Are QEs serving student and program needs?

The last two questions are critical as there remains a shared goal of promoting diversity in STEM PhD careers; furthermore, the job market for PhDs has greatly expanded since QEs were first accepted into the training canon meaning that training needs have evolved as well.

#### QE timing

Literature reports that QEs are offered sometime between the end of the first year or when courses are completed and the start of the fourth year, depending on several variables (Barnett et al., 2017; Bernstein et al., 2014). One variable that impacts timing is the institutional, doctoral program, or departmental policies and requirements. For example, Berkely’s PhD program in Mathematics QEs include a written preliminary exam before the end of the first three semesters and an oral QE that must be attempted by the end of year two (Earl▫Novell, 2006), one EdD program gives the QE in the spring semester of their second year, which is the last year of course work (McBrayer et al., 2020), marketing programs often have a first year QE and a second written QE due in year 2 or 3 (Ponder et al., 2004), and QEs are typically given during the second year of most nursing PhD programs (Allard et al., 2021). Other possible variables that may impact timing of a particular student’s completion of QEs include individual student preparedness, bureaucratic/administrative challenges like getting the assessment committee together, and lack of incentives to or penalties not to adhere to established timelines.

An important historical context of note is that the timing of QEs was designed by some programs to produce an early “off ramp” for doctoral students allowing students who are not prepared for or fully vested in completing dissertation research to exit the program be “weeded out” as it has been historically described. However, high attritions rates are not typically desired in modern doctoral programs, and hence programs generally seek to reduce or avoid inducing the potential student failure and stress with QEs that is associated with student attrition (Estrem and Lucas, 2003; McBrayer et al., 2020; Wilson et al., 2018). Interestingly, QE scores only weakly correlated with degree completion in one small study of an EdD program (McBrayer et al., 2020), and the timing of QEs did not negatively affect obtaining candidacy up to the eleventh semester in a broad study of PhD trainees (Ampaw and Jaeger, 2012). An intriguing idea is to customize the timing of QE administration based on student needs and programmatic outcomes in a way that considers competency-based learning. Related to this, a guide for competency-based assessments in biomedical PhD training has been proposed (Verderame et al., 2018).

#### QE format

Generally, QEs are written, oral, or a combination. Written QEs can include closed-book or open-book formats; timed or open-ended; and may include options such as on-site exams, take-home exams, written reports, and research proposals. Performance assessments are typically provided by groups of faculty, often in the form of an examining committee. Oral QEs can include presentations of a written proposal that can be based in the thesis proposed thesis research, presentation of selected topics (Mawn and Goldberg, 2012), or examination of student knowledge of critical thinking on select topics (Earl▫Novell, 2006).

For programs that use both written and oral QEs, they are sometimes considered distinct steps towards candidacy in terms of timing, name, and focus. For example, the written portion might be considered a comprehensive or preliminary exam and be based on coursework while the oral QE is based on proposed research or assigned papers. However, there is a lack of consistency in the distinctions between oral and written exams, including how they referred to by various programs. We found that some programs have explored alternative approaches. For example, some political sciences PhD programs provide students several options for QEs (Schafer and Giblin, 2008). In Counselor Education Doctorate programs, an alternative approach using student portfolios in place of comprehensive and final oral exams has been implemented (Cobia et al., 2005). A similar student-centered approach utilizing student-developed “authentic assessments” associated with standard and optional benchmarks has been implemented in a recently established EdD program in place of traditional QEs (Stewart▫Wells and Keenan, 2020).

Aligned with the idea that nontraditional QEs would provide greater value than closed-book written exams, students in engineering PhD students “showed interest in having the exam be focused on the research proposal as opposed to background theory and that they would appreciate more feedback and communication with the committee” (Memarian et al., 2019). One consistent theme from our review of the literature is that knowledge-based, written assessment is considered appropriate for time/location constrained closed book exams whereas open book, take-home written assessment and other alternative assessments, such portfolios, are more appropriate to assess higher order thinking (Kostohryz, 2016; Pelfrey and Hague, 2000; Ponder et al., 2004). One concern with alternative assessments for QEs is the challenge of standardizing process (Estrem and Lucas, 2003; Stewart ▫ Wells and Keenan, 2020). However, one common alternative is to include a thesis prospectus in the QE which has been reported to correlate with lower attrition and shorter time to completion (Nerad and Cerny, 1993).

#### Student experience with QEs

As a high-stakes milestone essential to the advancement to doctoral candidacy, the QE can be stressful and strategies to understand and enhance the student experience with QEs have been reported. This is especially important in the context of diversity and inclusivity as QEs have been identified as a potential “leak in the STEM pipeline” (Wilson et al., 2018). Vagueness in expectations and lack of understanding of the exam itself increases stress and anxiety significantly, which can impact performance (DiPietro et al., 2010; Harding▫DeKam et al., 2012; Nerad and Cerny, 1993). An older report found that students overprepared as a result of the lack of clarity, suggesting that making expectations, format, and other details associated with QEs clear will benefit students (Nerad and Cerny, 1993). Kostohyrz and colleagues reported that 22% of reviewed doctoral programs in counselor education had no written policy for evaluating questions (Kostohryz, 2016). Similarly, 97% of reviewed comprehension and rhetoric programs gave no public statement on how exams are assessed (Estrem and Lucas, 2003). Nursing doctoral students also voiced a need to better understand evaluation criteria and exam outcomes, as variable qualifying exam experiences created a feeling of injustice and during preparation for the QE students exhibit the poorest well-being (i.e., highest stress levels and lowest satisfaction with program) (Allard et al., 2021). The interviewed nursing students suggested that more support would de-stress the QE experience, such as senior student advice to help prepare for QEs as well as peer study groups.

One approach to overcome the stress associated with QE “unknowns” is to utilize an external mentoring model (Williams et al., 2017). In this model, social support from a national network, which students perceived as critical to effectively preparing for and dealing with the stress of QEs. The work of Williams and colleagues (2017) suggests that implementing a mentoring program can enhance the success of underrepresented groups on QEs and beyond, which is notable given the need to increase diversity of PhD researchers. Another support mechanism is to provide mock QEs for students; this approach was successfully implemented at the MD Anderson Graduate School of Biomedical Sciences to increase retention specifically of historically underrepresented minorities (Wilson et al., 2018). In addition to providing support and clarity, the format and flexibility of QEs also impact student perceptions. For example, Stewart Wells and colleagues found that students assessed using “authentic assessments” including writing research conference proposals or grant proposals and publishing an article in a peer-reviewed journal reported positive and motivating experiences (Stewart▫Wells and Keenan, 2020). Though limited evidence-based research has been completed in the biomedical sciences, broadly across disciplines, the evidence is clear: QEs are stressful and mechanisms to decrease stress and decrease QE associated attrition are all reported to benefit student experiences with QEs, including addition support, clarifying the QE purpose, format and assessment mechanisms, and finding ways to increase flexibility of QEs.

#### QE value and equity

Few studies have tackled complex questions regarding the equity of QEs in doctorate training, and we could not identify any studies that examine evidence-based associations between QEs and postgraduate outcomes. While program structure and support has been associated with attrition, the role of QE in attrition has been only briefly addressed. The most definitive studies on overall attrition in STEM fields are from the Council of Graduate Schools, which found that students starting STEM PhD programs between 1992-1993 and 2003-2004 had 46% completion rates at seven years and 57% at ten years for all students. A separate analysis showed that ten-year completion rates for students from URM background were slightly lower, 55% for White students, 51% for Hispanic students, and 47% for Black students, however the most recent CGS report on a larger set of URM students showed a 54% ten-year completion rate (Sowell et al., 2015), which suggests that there might not be a substantial demographic difference in overall completion rate.

However, attrition rates remain high, differ from program to program, and can in part be attributed to program environment and support. Likewise, an earlier study found similar attrition rates and determined that program culture and student fit were the most likely causes of attrition (Lovitts and Nelson, 2000). To lend further support to the thought that programs contribute to attrition, Ealr-Novel found higher rates of attrition for women and URM groups despite possessing similar GPAs (Earl▫Novell, 2006). It was hypothesized the higher attrition stemmed from a lack of mentorship and role models for these groups in PhD programs. Not surprisingly, one aspect of PhD programs that may have a potentially negative and highly influential impact on equity may be QEs. The high stakes and stress inducing QE process has been identified as a contributor to URM student attrition, while noting that program support can mitigate this problem (Wilson et al., 2018). It seems natural that QEs as traditionally implemented would lead to attrition, and the evidence suggests that providing support and potentially changing the format of QEs can lower attrition rates.

A separate question is whether QEs contribute to any bias in completion rates. While potential bias has been studied in programs as a whole, there is little literature exploring bias in the QEs. By contrast, a great deal of evidence has established bias in standardized exams such as the graduate record examination (GRE), which correlate with applicant demographics but not PhD program outcomes, such as first-author publication or time-to-degree completion, suggesting that these exams might inappropriately limit admittance of minority groups that historically perform less well on the exam (Hall et al., 2017; Moneta▫Koehler et al., 2017). As a result, many programs have explored dropping GRE requirements. Hence, following a similar argument, if it were found that QEs in fact introduced bias in program advancement and attrition, they could be inadvertently harming efforts to increase diversity of the STEM PhD workforce. It is important for programs to consider any potential biases introduced by the QE process, and proactively establish mechanisms to mitigate bias and ensure equity in the QE process.

Our review of the literature shows that there is little consistency across doctorate programs on QE format, assessment, timing, student support, or even nomenclature. QEs appear to contribute to attrition rates and increase student stress. For STEM PhD programs specifically, we did not find substantial evidence supporting any specific QE practices except for providing clarity to students of QE purpose and format and providing student support during QEs.

### QUALIFYING EXAMS: ANALYSIS OF PEER UNIVERSITY WEBSITES

The literature survey revealed mostly descriptive research on QEs, with a notable absence of QE roles in biomedical, pharmaceutical sciences, and adjacent PhD programs. To begin the process of filling in this critical gap in our understanding of QE practices in PhD training, we examined the websites of 23 universities: 22 peer universities (i.e., University- and School-defined peers) and internal university-wide policies institutionally on QEs. We specifically evaluated the prerequisites, timing, nature, established term for QEs, as well as outcomes and objectives stated for QEs. To our knowledge, this represents the first analysis of central regulations regarding QEs representative of major U.S. Research intensive universities.

#### General Characteristics of University-wide QE Policies

Of the 23 reviewed Universities (Table 1), most (n=20) have a specific Graduate School or College that sets policies for graduate education, while three have graduate administration delegated to discipline-specific Schools/Colleges. Two of the Universities without a Graduate School/College maintained central oversight of graduate education through the provost office that was guided by a faculty committee. Of the 20 Universities with a central Graduate School, nearly all (n=19) explicitly state a QE requirement for the PhD. One University without a Graduate School stated a university-wide QE requirement. Of the 3 Universities without a specific central QE requirement, it is stated to be or assumed to be left to individual PhD programs, and each biomedical program examined at these Universities indeed required some form of QE.

**Table 1.** Characteristics of Qualifying Exam (QE) Policies for UNC and Peer Institutions (N = 23)^1^.

| Research Question | Yes<sup>2</sup><br>n(%) | Summary of Website Data Collected |
| --- | --- | --- |
| **Does university have a Graduate School or College?** | 20 (87%) | Two of the Universities without a graduate school oversee graduate programs through the provost's office with a faculty committee providing guidance. The other University delegates oversight to individual Schools and Colleges without any indication of central oversight. |
| **Does university have a QE requirement?** | 20 (87%) | Of the 20 "yes" Universities, 19 state QE requirements of the graduate schools. Universities without a requirement leave it to individual programs, and all biomedical sciences programs checked required a QE. |
| **Question asked for universities with a Graduate School/College (n=20)** |  |  |
| **Is it called a Qualifying Exam?** | 7 (35%) | Other descriptors include Preliminary Exam (n=9), Doctoral Exam (n=2), Doctoral Comprehensive Exam (n=1), Comprehensive Exam (n=1), Candidacy Exam (n=1), and General Exam (n=1) |
| **Is passing the QE required to advance in the program?** | 20 (100%) | Policies for retaking the QE varies from school to school (in case of failure) |
| **Does student status change after passing the QE?** | 19 (95%) | Eighteen call the students "candidates", one calls them ABD (All But Dissertation). One provided a certificate, and one held a ceremony. |
| **Do students need to be registered for QE?** | 15 (75%) |  |
| **Is a thesis proposal/proposal required?** | 9 (45%) | The prospectus can be part of the QE. For each, a proposal or prospectus is commonly part of qualifying exam practices even though not explicitly required. |
| **Is the QE format described?** | 8 (40%) | Two use oral format, six use oral and written format |
| **Is a time limit for QE passing described?** | 7 (35%) | Programs require QE completion within 3 years, 4 years, or 5 years |

<sup>1</sup> We explored the publicly accessible websites of the universities to evaluate campus-wide policies on QEs:
<sup>2</sup> Yes indicates that the Graduate School or campus-wide resource site explicitly described or stated a given policy.

We found that all but one University that required a QE advanced the PhD students’ status upon passing to “Candidate” after passing the QE. Passing QEs were commemorated as well with one University providing a “Candidate Certificate” and another holding a ceremony to commemorate obtaining candidate status. Since QEs were considered a requirement for the PhD in most programs, we anticipated that Universities would have a registration requirement and found that 15 of the 20 Graduate Schools/Colleges did have a requirement requiring students to be actively registered in for-credit coursework when the QE was taken. We did not find any Universities that offered/required a specific QE course. We also explored whether a thesis proposal/prospectus was required as part of the QE, however we instead concluded that while it was a common option, it was not typically a requirement. We found evidence that several graduate schools (n=9) generally required a thesis proposal/prospectus. Each of these graduate schools suggested that the proposal/prospectus could be considered for the QE, but none of the Graduate Schools *required* that the proposal/prospectus specially be part of the QE.

Graduate Schools referred to QEs as preliminary exams (n=9), qualifying exams (n=7), candidacy exams (n=1), and General Exam (n=1). Irrespective of nomenclature, all graduate Schools clearly defined QEs as a major milestone and 18 (90%) defined this achievement as the primary prerequisite for candidacy status. The format and timing of the QE also varied across Schools. Of those that described the QE format (n=8, 40%), 10% of those (n=2) use oral formats and 30% of those (n=6) use some combination of oral and written formats. The other programs (n=12, 60%) either did not describe the QE format, or indicated that the format was determined by each program. In several programs, 45% (n=9), a prospectus of the proposed thesis research was explicitly required, and the Graduate School allows programs to use the prospectus as the written QE. We did not find evidence for any further regulations on the QE format, for example in class/take home, open/close book, course review, literature review. Similarly, we did not find any evidence regulating the specific nature of the oral exam, and it was apparent that these details were left to individual programs to define. Timing of QEs was explicitly regulated by some graduate schools (n=7), with these schools requiring the exam within 3, 4, or 5 years of enrolling in the program. Other timing regulations we observed included schools that set the timing of the QE as before (the quarter or three months) the final defense (n=4) and some schools stated that the defense had to be within 5 years after the candidacy exam or QE must be retaken (n=3). Several graduate schools (n=) also set requirements prior to allowing students to sit for the QEs such as certain number of credit hours, completing required courses, and being in good standing as defined by no incomplete or failing grades.

Because QEs are a major gatekeeper of PhD student progression, it is critical to consider policies QEs assessment, opportunities for remediation, and program alignment. Each graduate school stated that QEs were to be assessed by a faculty committee. Graduate schools varied on whether the QE committee and dissertation committee were the same, with 5 schools stating the committees are the same and 2 stating they are different but can overlap. Other schools appear to leave this to the individual PhD programs. QE outcomes were established via the following common committee voting norms: as unanimous decision of committee (n = 3), only up to one “failure” vote (n = 5), and majority vote to pass (n = 4). For the other schools (n = 4), it was left to the individual programs to set the policy to determine passing. Generally, the QE committee also determines if a retake will be allowed, and all schools stated that an opportunity for remediation should be considered with a variety of school-specific policies regulating a possible retake.

Regarding program alignment, several schools (n = 11) indicated a relationship of the QE to program outcomes. There was a broad range of details in these sections with some descriptions being quite general for example indicating the QE should “provide written evidence of competency in the field” and others providing a more comprehensive description of the intent of the QE by expanding on specific competencies, for example mastery of the literature and critical thinking, expected of PhD trainees. Only one school indicated that programs must “Establish and communicate the program-specific purpose, format, and evaluation of the doctoral preliminary written examination” and provides a guide to help programs evaluate the QE for program alignment.

## CONCLUSIONS & FUTURE DIRECTIONS

It is clear from our review of the literature and peer websites that QE exams have a wide variety of names, formats, and regulations. Despite these variances, there appears a shared overarching purpose: a high-stakes assessment that doctoral students complete as a prerequisite to fully engaging independent dissertation research. In our evaluation, we identified some QE processes that evidence suggests will enhance student outcomes. Specifically, creating transparency in QE practices with explicit expectations and building structured student support are steps programs can take to improve QE outcomes and we call for more research to help establish evidence-based best practices in QEs for doctoral training.

Future work should develop best practices for QE format and assessment as well as evaluate the relationship between QEs and student outcomes. In addition, the functional purposes that QEs serve should be better explored to better understand whether the QE evaluation methods are in fact predictive of any outcomes and future professional success. We propose that addressing the following topics as the basis of future studies and for programs to consider in evaluating their approach to QEs:

- Identify faculty, students, and administrator perceptions regarding current and best QE practices
- Assess QEs alignment with program competencies
- Assess QEs alignment with program needs
- Assess alignment with QEs student needs
- Evaluate QE practices for potential bias and identify strategies for mitigation
- Evaluate effectiveness of specific QE formats
- Evaluate the impact of QE format and practices on student outcomes

QEs serve as an important milestone, programmatic element, and (unfortunately, at times) a gatekeeper to progress to advanced stages of training. By addressing these important issues with evidence-based policies and practices, there is the potential to create more effective evaluation tools, align associated outcomes with training goals, improve the student experience, and create a training culture and research enterprise with greater equity. While these may seem like lofty goals, effectively preparing the future biomedical workforce is worth the effort, and an empirical approach to QEs will go a long way toward achieving progress across these priorities.

## ACKNOWLEDGMENTS

This work was supported in part by National Institutes of Health General Medical Sciences - Science of Science Policy Approach to Analyzing and Innovating the Biomedical Research Enterprise (SCISIPBIO) Award (GM-19-011) - 1R01GM140282-01.

